# Wnt signalling is a major determinant of neuroblastoma cell lineages

**DOI:** 10.1101/506980

**Authors:** Marianna Szemes, Alexander Greenhough, Karim Malik

## Abstract

The neural crest, which has been referred to as the fourth germ layer, comprises a multipotent cell population which will specify diverse cells and tissues, including craniofacial cartilage and bones, melanocytes, the adrenal medulla and the peripheral nervous system. These cell fates are known to be determined by gene regulatory networks (GRNs) acting at various stages of neural crest development, such as induction, specification, and migration. Although transcription factor hierarchies and some of their interplay with morphogenetic signalling pathways have been characterised, the full complexity of activities required for regulated development remains uncharted. Deregulation of these pathways may contribute to tumourigenesis, as in the case of neuroblastoma, a frequently lethal embryonic cancer thought to arise from the sympathoadrenal lineage of the neural crest.

In this conceptual analysis, we utilise next generation sequencing data from neuroblastoma cells and tumours to evaluate the possible influences of Wnt signalling on neural crest GRNs and on neuroblastoma cell lineages. We provide evidence that Wnt signalling is a major determinant of regulatory networks that underlie mesenchymal/NCC-like cell identities through PRRX1 and YAP/TAZ transcription factors. Furthermore, Wnt may also co-operate with Hedgehog signalling in driving proneural differentiation programmes along the adrenergic lineage. We propose that elucidation of Signalling Regulatory Networks can augment and complement GRNs in characterising cell identities, which will in turn contribute to the design of improved therapeutics tailored to primary and relapsing neuroblastoma.

## Introduction

Neuroblastoma (NB) is a frequently lethal paediatric tumour, with 75% of NBs occurring in children under 5 years of age. About half of the tumours arise in the adrenal medulla, with the remainder originating in the paraspinal sympathetic ganglia in the abdomen or chest, or in pelvic ganglia. This distribution reflects the probable developmental origin of NB in the sympathoadrenal (SA) lineage of the neural crest (NC). Clinically, NB encompasses low-risk disease which responds well to treatment and may even spontaneously regress, and high-risk disease, representing about 40 % of total cases, which frequently relapse and have less than 50 % survival. This clinical heterogeneity is thought to reflect both the complex molecular etiology as well as the cellular heterogeneity of tumours (Brodeur 2003; Maris et al. 2007). Whilst it has long been known that the protooncogene *MYCN* is crucial for neural crest cell fate (Wakamatsu et al. 1997) and influencing differentiation states in NB (Westermark et al. 2011), a deeper understanding of developmental factors remains necessary to determine the origins of NBin order to inform improved prognosis and therapies.

In this conceptual analysis, we will consider the potential regulatory influences and interactions of the canonical Wnt signalling pathway in determining phenotypes in the neural crest and neuroblastoma. Although this signalling pathway is known to be critical in regulating stemness, cell fate, differentiation and proliferation (Nusse and Clevers 2017), much remains unclear about its role in neuroblastoma. Based on our recent identification of genes regulated by the Wnt ligand Wnt3a and Wnt agonist R-spondin 2 in a NB cell-line (Szemes et al. 2018), we assess transcriptional and signalling pathways which require further investigation in the contexts of neural crest development and neuroblastoma.

### Wnt signalling pathways and components

In broad terms, Wnt signalling includes “canonical” and “non-canonical” pathways. The former is also referred to as Wnt/β-catenin signalling, as it is dependent on cytoplasmic-nuclear translocation of β-catenin, and its subsequent transcriptional cofactor activity with T-cell factor/lymphoid enhancer factor (TCF/LEF) transcription factors (Clevers and Nusse 2012; Nusse and Clevers 2017). Non-canonical, or alternative Wnt signalling pathways include the Planar cell polarity (PCP), the Wnt/Ca^2^+ pathways, β-catenin-independent pathways acting via Rho-associated kinase (ROCK) and G-protein dependent calcium release respectively (Komiya and Habas 2008). More recently, another alternative Wnt pathway has been demonstrated, where the downstream effectors are the Hippo signalling pathway transcriptional co-factors, Yes-associated protein (YAP) and Transcriptional C-activator with PDZ-Binding motif (TAZ, encoded by *WWTR1)* (Park et al. 2015) (Figure 1).

**Figure 1.**
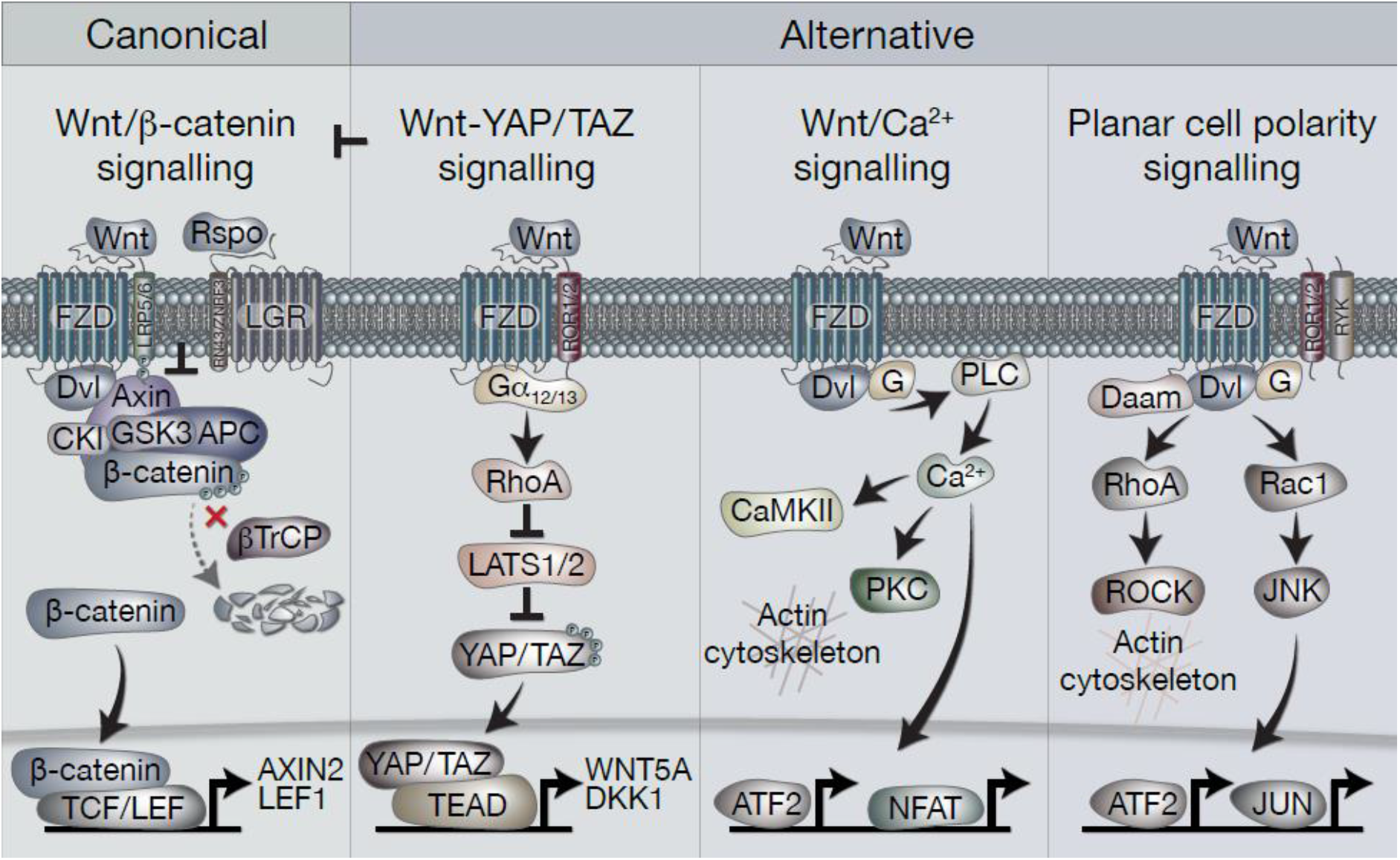
Overview of canonical and alternative Wnt signalling pathways. **Canonical Wnt signalling** is defined by the activation of β-catenin-dependent transcription (via TCF/LEF) downstream of Wnt receptors. In the absence of Wnt ligands, β-catenin is held in the destruction complex of proteins that includes Axin, APC, Ser/Thr kinases CK1 and GSK3, and E3-ubiquitin ligase β-TrCP. Sequential phosphorylation (by CK1 and GSK3) and ubiquitination (by β-TrCP) of β-catenin promotes its proteosomal degradation. Binding of Wnt ligands (classically Wnt3a) to FZD and LRP5/6 receptors leads to recruitment of the destruction complex to the membrane via Dvl and Axin, which blocks β-catenin ubiquitination by β-TrCP. The destruction complex then becomes saturated with β-catenin, allowing newly synthesised β-catenin to accumulate in the cytoplasm and translocate to the nucleus where it associates with TCF/LEF transcription factors to regulate gene expression. Rspo (R-spondin) proteins can enhance Wnt signalling by binding LGR4/5/6 receptors to antagonise the RNF43/ZNRF3 transmembrane E3 ligases that remove Wnt receptors from the cell surface. **The alternative Wnt-YAP/TAZ pathway (as defined by Park et al 2015).** Wnt5a/b and Wnt3a ligands induce the activation of YAP/TAZ via FZD and ROR1/2 co-receptors, independently of LRP5/6 and β-catenin. Activation of FZD-ROR1/2 couples to Gα12/13 G-protein subunits, leading to activation of RhoA and subsequent inhibition of LATS1/2 kinases (major YAP/TAZ Ser/Thr kinases). Inhibition of LATS1/2 leads to YAP/TAZ dephosphorylation, stabilisation and translocation to the nucleus where their interaction with TEAD transcription factors promotes gene regulation. Wnt-YAP/TAZ target genes such as DKK1 lead to the inhibition of canonical Wnt/β-catenin signalling. **Wnt/Ca2+ signalling.** Activation of FZD stimulates the activity of PLC (phospholipase C) via Dvl and G proteins, leading to increases in intracellular Ca2+ levels. Ca2+ activates CaMKII (calmodulin-dependent protein kinase II), PKC (protein kinase C) and the transcription factors NFAT and ATF2. Activation of PKC by Ca2+ can cause actin cytoskeleton rearrangements via Cdc42 activity. **Wnt/PCP (planar cell polarity) signalling.** Wnt signalling via FZD with co-receptors ROR1/2 or RYK may activate G proteins and Dvl to stimulate RhoA (via Dvl-Daam) and ROCK activity, leading to rearrangement of the actin cytoskeleton. Activation of G proteins can also stimulate Rac1, leading to JNK-mediated phosphorylation of c-JUN and transcriptional activity via AP-1.

There is complex and extensive interplay between these pathways, reflected in the 19 Wnt ligands and 10 Frizzled (FZD) receptors. Wnt ligands are secreted, lipid modified glycoproteins conserved in all metazoan animals, able to act as morphogens over short distances (MacDonald, Tamai, and He 2009). Although some preferential usage of certain Wnt ligands by different pathways is observed, such as Wnt3a for canonical, and Wnt5a for non-canonical signalling, strict restriction of ligand usage by pathways, or a “Wnt code”, have not emerged. Rather, numerous other effectors and regulatory molecules add to the complexity of the pathways (van Amerongen 2012). In general, however, non-canonical Wnt signalling is thought to antagonise the canonical Wnt/β-catenin pathway (Veeman, Axelrod, and Moon 2003).

In the canonical Wnt/β-catenin pathway, Wnt ligands are bound with high affinity by Frizzled proteins, leading to formation of heterodimeric membrane core receptor complexes containing LRP5/6. Binding of Wnt ligands leads to LRP phosphorylation and recruitment of the scaffold protein Axin to LRP. Axin is a key component of the “destruction complex”, which also includes APC, the Ser/Thr kinases GSK-3 and CK1, protein phosphatase 2A (PP2A), and the E3-ubiquitin ligase β-TrCP. This complex regulates cytoplasmic β-catenin turnover through phosphorylation and proteosomal turnover. The LRP-Axin interaction leads to inactivation of the destruction complex, permitting β-catenin stabilization and nuclear availability. The TCF/LEF family of transcription factors are then able to utilise β-catenin as a transcriptional co-activator, and instigate target gene expression (Clevers and Nusse 2012). Further amplification of signalling can be achieved through the participation of another set of receptors, the leucine-rich repeat-containing G-protein coupled receptors (LGR4/5/6) and their ligands, the R-spondins (Rspos) (de Lau et al. 2011). LGR-Rspo complexes at the cell membrane decrease the endocytic turnover of Frizzled-LRP5/6 by neutralising the ubiquitin ligases RNF43 and ZNRF3 (Hao et al. 2012).

### Differentiation of the neural crest and Wnt signalling

The tissue of origin for neuroblastoma is the neural crest, a band of cells that forms transiently between the neural tube and the non-neural ectoderm during early vertebrate development. Many signalling pathways act together during gastrulation to promote neural crest induction and specification, especially bone morphogenetic protein (BMP) (Steventon et al. 2009), fibroblast growth factor (FGF) (Stuhlmiller and Garcia-Castro 2012) and Wnt/β-catenin signalling (Leung et al. 2016; Garcia-Castro, Marcelle, and Bronner-Fraser 2002). Together, these signalling pathways orchestrate numerous transcription factors, including MSX1/2 (Ramos and Robert 2005; Tribulo et al. 2003) and ETS1 (Barembaum and Bronner 2013). Neural crest specification finishes with elevation of the neural folds during neurulation, and thereafter, following neural tube closure, premigratory neural crest cells delaminate and undergo epithelial-mesenchymal transition (EMT) (Nieto et al. 2016). Multipotent NCCs with differential abilities to form derivative cell types are arranged along the length of the vertebrate embryo; these can be subdivided into the cranial, vagal, trunk and sacral neural crest, and will give rise to diverse cell and tissue types, including facial cartilage and bone, melanocytes and smooth muscle (Martik and Bronner 2017; Bronner and LeDouarin 2012). The trunk NCCs give rise to the sympathetic nervous system and the adrenal medulla and are therefore the source of presumptive progenitors of neuroblastoma cells.

As well as being crucial for neural crest induction, canonical Wnt signalling has also been shown to be vital in other NC stages including a role in delamination in co-operation with BMP; here disruption of β-catenin and TCF/LEF inhibits cell cycle progression and neural crest delamination and transcription of BMP-regulated genes, including *MSX1* (Burstyn-Cohen et al. 2004). Furthermore, ablating β-catenin specifically in neural crest stem cells *in vivo* revealed that, despite some neural crest–derived structures developing normally, mutant animals lack melanocytes and dorsal root ganglia. β-catenin mutant NCCs appear to emigrate normally but fail to undertake sensory neurogenesis, thereby suggesting a role of β-catenin in premigratory/early migratory NCCs (Hari et al. 2002). Non-canonical Wnt signalling is also involved in NCC migration (Mayor and Theveneau 2014). Thus, Wnt signalling pathways are involved at multiple stages of NC development and can also selectively regulate NC lineages.

The aforementioned signalling pathways orchestrate cell behaviour via gene regulatory networks (GRNs) that govern neural crest development. Transcriptional analysis and molecular loss and gain-of-function studies facilitate a GRN model of the NC, which, encompassing transcriptional regulators and signalling molecules, can explain the formation and maintenance of NC lineages (Martik and Bronner 2017). Establishment of the neural plate border, specification, migration and differentiation, have their specific GRNs, and these networks can interact, overlap and influence each other. Gradients and balances of Wnt, BMP, FGF and Notch signalling determine NC induction and activate transcription factors (e.g. *MSX1, MYCN, DLX5/6*), in the neural plate border module, determining the boundaries of NC, neural and non-neural ectoderm. These in turn, upregulate NC specifier genes, which initiate the EMT programme (including *SNAI1/2, ETS1, TWIST1/2*) to prepare for delamination of NCCs as well as maintain an undifferentiated state (premigratory module). Following EMT, the NCCs acquire a migratory programme enabling long-distance migration guided by environmental signals and maintained by a TF network (including ZEB2, LMO4, SNAI1/2, TWIST1/2, ETS1). The sympathoadrenergic lineage follow a ventral migratory pattern and aggregate at the dorsal aorta and go on to form the primary sympathetic ganglia and colonize the adrenal medulla. BMPs released by the wall of the dorsal aorta induce SA speciation, activating *ASCL1* and *PHOX2B*, which switches on other SA-lineage specific transcription factors, for example ASCL1 upregulates *PHOX2A*, which in turn upregulates *TH* and *DBH*, enzymes characteristic of the SA lineage.

*MYCN*, the key developmental transcription factor deregulated in neuroblastoma, has been shown to be a Wnt target gene in chicken limb mesenchyme (ten Berge et al. 2008). *MYCN* is expressed at high levels in chick neural crest progenitors in early NC development, but surprisingly is not apparent in highly proliferative migrating neural crest cells (Khudyakov and Bronner-Fraser 2009). This finding was confirmed and extended to show absence of *MYCN* expression in the condensed dorsal root and sympathetic ganglia. Elevating *MYCN* expression in the neural plate border was shown to lead to a change in neural crest identity, toward a more central nervous system fate (Kerosuo et al. 2018). Transient overexpression of ectopic *MYCN* in early migrating cells, however, suggested that MYCN can regulate neural crest fate by regulating ventral migration and neuronal differentiation (Wakamatsu et al. 1997). Evidence from the developing NC in lamprey suggests that ZIC, MSX and TFAP2A/AP-2 transcription factors may positively regulate *MYCN* in the NC (Nikitina, Sauka-Spengler, and Bronner-Fraser 2008), but signalling pathways upstream of *MYCN* in the NC are unclear.

### Neuroblastoma and Wnt signalling

Neuroblastomas display relatively few gene mutations, thus limiting the options for targeted therapeutics, and relapsing disease is common for high risk NB (Matthay et al. 2016). Another increasingly important consideration is cellular heterogeneity of NBs. Although this was first observed decades ago in NB cell lines being able to transdifferentiate (Ciccarone et al. 1989), the dependence of lineage identity on transcriptional circuitries are only recently becoming evident. One study recently established that primary NBs and NB cell-lines can contain two major cellular components, including migrating, neural crest cell-like mesenchymal (MES), and more committed adrenergic (ADRN) cells. These two cell types were defined by super-enhancer-associated gene expression patterns, characteristic of neural crest lineage differentiation stages and could transdifferentiate *in vitro* and *in vivo*. The ADRN lineage was defined by transcription factors such as GATA2/3, PHOX2A/2B and DLK1, whereas the MES lineage included the classical EMT transcription factor SNAI2, together with YAP1, TAZ (encoded by *WWTR1)* and PRRX1. The ADRN cells were more tumorigenic in nude mice than MES cells, although MES cells were more chemoresistant and were enriched in relapsed tumours (van Groningen et al. 2017). Concomitantly, another group also established transcriptional circuitries underlying the heterogenous differentiation states of NB cells, and termed their 3 subtypes as sympathetic noradrenergic, NCC-like and a mixed type (Boeva et al. 2017). The noradrenergic identity required a transcription factor module containing PHOX2B, HAND2, and GATA3 and therefore corresponds to the ADRN subtype defined by van Groningen *et al* (van Groningen et al. 2017), whereas the NCC-like overlaps with the MES subtype, given they both require PRRX1. Importantly, ectopic overexpression of PRRX1 was shown to be able to alter lineage identity from ADRN to MES cells (van Groningen et al. 2017).

The prototypic route for disrupted differentiation of NCCs leading to neuroblastoma depends on *MYCN* amplification (Brodeur et al. 1984), which in turn leads to overexpressed MYCN protein directly repressing genes required for sympathetic nervous system terminal differentiation (Westermark et al. 2011; Gherardi et al. 2013). Given the many examples of cancers dependent on oncogenic Wnt/β-catenin signalling (Clevers and Nusse 2012), such as colorectal cancers which have activating mutations in β-catenin or loss of function APC mutations (Fodde and Tomlinson 2010), and that *MYCN* is a target of Wnt signalling (ten Berge et al. 2008), it was reasonable to propose that Wnt/β-catenin signalling would represent an oncogenic pathway in NB. This was further supported by our demonstration that high levels of LGR5 were apparent in undifferentiated NBs and NB cell-lines (Vieira et al. 2015) and *LGR5* expression correlated with poor prognosis. However, although we showed that LGR5 was able to function as a Rspo2 receptor and amplify Wnt3a-induced β-catenin/TCF transcriptional activity, marked increases in proliferation were not observed, and the pro-survival functions of LGR5 were shown to be attributable to LGR5 positively modulating MEK/ERK signalling rather than Wnt/β-catenin signalling (Vieira et al. 2015). This dual regulatory capacity of LGRs on MEK/ERK signalling and Wnt was subsequently also demonstrated in skin carcinogenesis (Xu et al. 2016), but the underlying mechanisms remain unclear.

Although we found that Wnt3a/Rspo2 treatment of NB lines lead to induction of some established canonical Wnt target genes, such as *LEF1* and *AXIN2, MYCN* transcripts were not induced. In fact, we found that MYCN and MYC protein levels were actually reduced after Wnt3a/Rspo2 treatment, in contrast to previous reports suggesting induction of *MYC* in non-*MYCN* amplified (non-MNA) NBs as a result of Wnt/β-catenin signalling (Liu et al. 2008). Furthermore, our study did not align simply with reports showing that individual Wnt pathway components were associated with chemoresistance (*FZD1*) (Flahaut et al. 2009), tumorigenic stem-like cells in human and mouse neuroblastoma (*FZD6*) (Cantilena et al. 2011), and increased NB proliferation dependent on FZD2 (Zins et al. 2016). Using Wnt chemical agonists and inhibitors another study suggested that Wnt signalling hyperactivation promotes apoptosis of NB cells, and that Wnt inhibition decreased proliferation and increased NB differentiation (Duffy et al. 2016). Oncogenicity of Wnt/β-catenin signalling, via deregulation of MYCN, was also inferred in a report demonstrating the therapeutic benefit of glypican-2 immunotherapy for NB (Li et al. 2017).

Given the lack of consensus on the role of Wnt signalling in NB, we therefore sought to detail the phenotypic and transcriptomic effects of Wnt signalling on NB cells by treating three cell lines with Wnt3a and Rspo2. We determined early (6 hrs) Wnt-induced transcriptomic changes by RNA sequencing in SK-N-BE(2)-C, a MYCN-amplified (MNA) NB cell line (Szemes et al. 2018). Using a statistical cut-off of p=<0.005, we identified 90 genes that showed substantial and significant changes. In parallel, longer treatments were assessed for changes in phenotype, and established that Wnt3a/Rspo2 treatment could induce proliferation in the SK-N-AS cell line, but in other lines, especially SK-N-BE(2)-C and SH-SY5Y cell-lines, increased proliferation was not observed. Instead we found an EMT-like transition, evidenced by increased cell migration and induction of *SNAI1/2* and *TWIST1/2*, and neural differentiation accompanied by increased expression of marker genes such as *NGFR, NTRK1, NEFM* and *NEFL/NF68*. Meta-analysis of the expression of our 90 Wnt target genes in primary tumour gene expression databases revealed (i) four gene subsets within the 90 genes, and (ii) dramatically different prognostic outcomes associated with each of the 4 gene subsets (Figure 2). By converting the expression of each of the four gene subsets into a numerical model, or metagene, we were further able to show that, for example, expression of Wnt metagene 1 (WMG-1) strongly positively correlated with Hippo-YAP/TAZ signalling signatures (R = >0.76) in NB, and that these genes were more highly expressed in intermediate risk NB, rather than low- or high-risk NB. WMG-3 was overexpressed in the high risk NB cluster, dominated by MNA tumours. Conversely, WMG-2 showed a strong positive correlation to Hedgehog signalling (R = >0.84) associated with differentiation (Souzaki et al. 2010) and a negative correlation with MYC signatures (R = - 0.76). WMG-2 genes were, concordantly, strongly repressed in high-risk MNA NB (Szemes et al. 2018).

**Figure 2.**
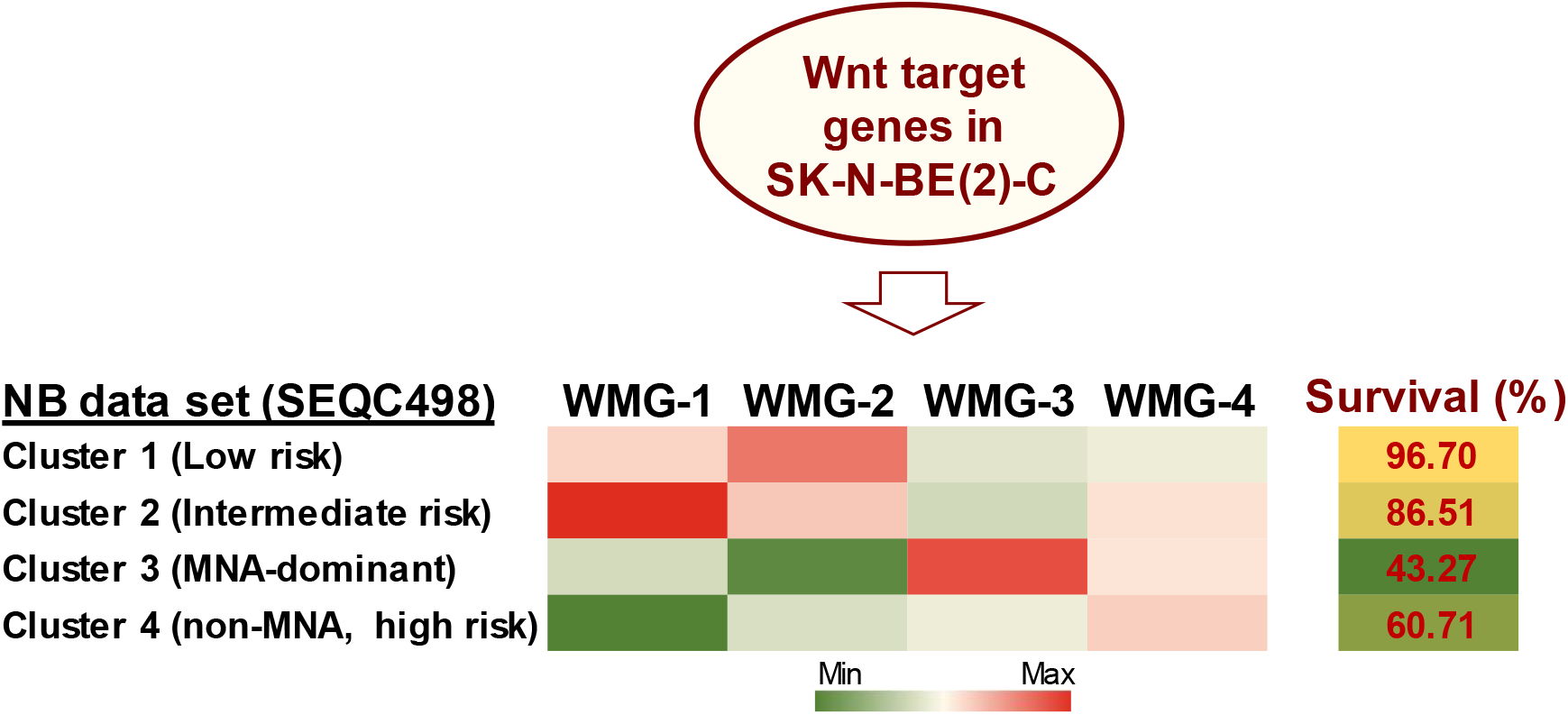
Summary of Wnt target genes previously identified in SK-N-BE(2)-C cell line. (Szemes et al. 2018). K-means clustering of these target genes in the SEQC NB dataset (GSE62564) enabled identification of 4 co-expressed Wnt target gene subsets. The combined expression of each gene subset was converted into a single value, represented as a metagene. The <u>W</u>nt <u>m</u>eta<u>g</u>ene groups (WMG-1,2,3,4) displayed differential expression in 4 prognostic clusters in the primary NB tumour data set. Expression levels of the WMGs in the NB clusters are indicated by the heatmap, and survival for each NB patient cluster is shown in the last column. Bioinformatic analyses were performed by using tools implemented in R2 (http://r2.amc.nl http://r2platform.com).

Collectively, studies from our laboratory and from others allude to an underappreciated role for Wnt signalling in NC and NB biology. In particular, the induction of EMT-like changes and the correlation of a subset of our Wnt target genes (WMG-1) with Hippo-YAP/TAZ pathway gene signatures, seemingly characteristic of the MES lineage (van Groningen et al. 2017), suggest that Wnt signalling may be part of the regulatory network determining NB cell identity. In the following sections, we will evaluate this concept through analysis of Wnt signalling on transcriptional and signalling pathways involved in NC and NB.

### Is Wnt signalling a major determinant of NC and NB cell fate?

Numerous studies have examined the spatiotemporal expression of transcription factors and signalling pathway components in the neural crest. Integration of these efforts has facilitated the establishment of a Gene Regulatory Network (GRN) underlying neural crest development (Martik and Bronner 2017). As our RNA sequencing of the NB cell line SK-N-BE(2)-C treated with Wnt3a/Rspo2 provides a unique gene set representing, at least in part, Wnt targets in the neural crest, we assessed which GRN genes encoding transcription factors or signalling molecules from different stages of NC development might be affected by Wnt3a/Rspo2 treatment (Table 1). As expected from our previous work, premigratory and migratory modules, including several EMT transcription factors (*TWIST1, ZEB2, ETS1*), are affected. Interestingly, given that the presumptive cell of origin for NBs is along the sympathoadrenal lineage, there is also strong induction for neural plate border specifiers *MSX1, DLX5* and *DLX6*. This suggests that either SK-N-BE(2)-C cells have retained early transcriptional plasticity, or more intriguingly, that NBs may arise prior to sympathoadrenal differentiation. This notion is supported by developmental experiments showing ectopic *MYCN* expression in the neural plate border resulting in a shift towards a CNS-like state (Kerosuo et al. 2018) and the induction of NB in MYCN-overexpressing NCCs (Olsen et al. 2017).

**Table 1.**
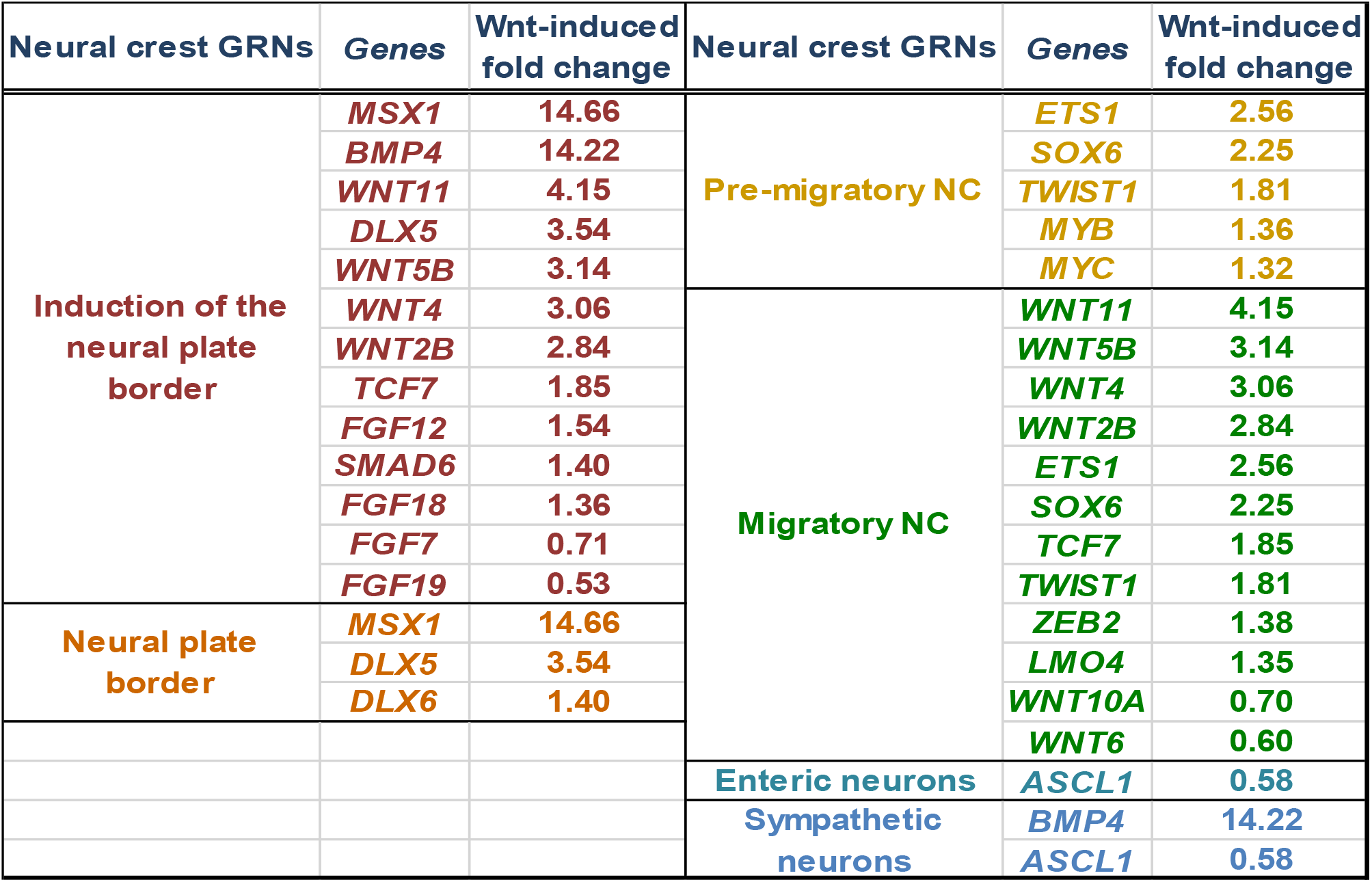
Effects of Wnt3a/RSPO-2 induction in SK-N-BE(2)-C neuroblastoma cells (ERP023744) on the gene regulatory network modules of the developing neural crest (as identified in (Martik and Bronner 2017). Regulatory genes with a minimum change of +/-1.5-fold are shown.

| Neural crest GRNs | Genes | Wnt-induced fold change | Neural crest GRNs | Genes | Wnt-induced fold change |
| --- | --- | --- | --- | --- | --- |
| Induction of the neural plate border | *MSX1* | 14.66 | Pre-migratory NC | *ETS1* | 2.56 |
|  | *BMP4* | 14.22 |  | *SOX6* | 2.25 |
|  | *WNT11* | 4.15 |  | *TWIST1* | 1.81 |
|  | *DLX5* | 3.54 |  | *MYB* | 1.36 |
|  | *WNT5B* | 3.14 |  | *MYC* | 1.32 |
|  | *WNT4* | 3.06 | Migratory NC | *WNT11* | 4.15 |
|  | *WNT2B* | 2.84 |  | *WNT5B* | 3.14 |
|  | *TCF7* | 1.85 |  | *WNT4* | 3.06 |
|  | *FGF12* | 1.54 |  | *WNT2B* | 2.84 |
|  | *SMAD6* | 1.40 |  | *ETS1* | 2.56 |
|  | *FGF18* | 1.36 |  | *SOX6* | 2.25 |
|  | *FGF7* | 0.71 |  | *TCF7* | 1.85 |
|  | *FGF19* | 0.53 |  | *TWIST1* | 1.81 |
| Neural plate border | *MSX1* | 14.66 |  | *ZEB2* | 1.38 |
|  | *DLX5* | 3.54 |  | *LMO4* | 1.35 |
|  | *DLX6* | 1.40 |  | *WNT10A* | 0.70 |
|  |  |  |  | *WNT6* | 0.60 |
|  |  |  | Enteric neurons | *ASCL1* | 0.58 |
|  |  |  | Sympathetic neurons | *BMP4* | 14.22 |
|  |  |  |  | *ASCL1* | 0.58 |

We next assessed the relationships between the Wnt pathway and neural crest cells and ARDN and MES identities using publicly available gene expression datasets. As shown in Figure 3A, transcription factors in the Wnt pathway are significantly higher in NCCs and MES cell lines compared with ADRN cell lines.

**Figure 3.**
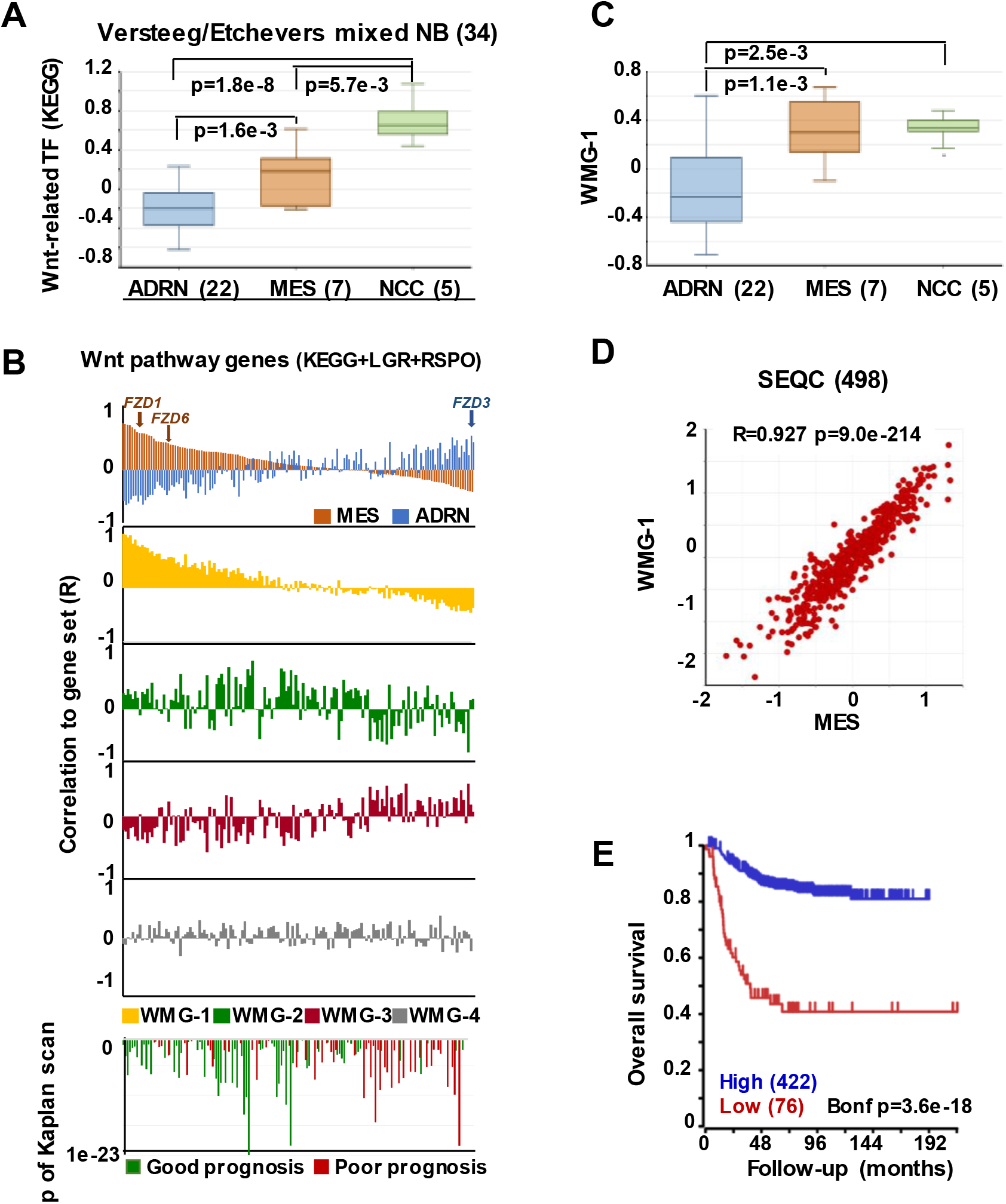
Wnt pathway associations with MES phenotype in neural crest cells and NB. **(A)** Expression of Wnt-related transcription factors (KEGG) in adrenergic (ADRN), mesenchymal (MES), and neural crest cell lines (NCC). **(B)** Correlation of the expression of Wnt pathway members (KEGG, LGRs and RSPOs; n=147) with various gene sets and prognosis in the SEQC data set (GSE62564), sorted according to the coefficient of correlation to MES signature. The x-axis represents Wnt pathway genes, while the y-axis indicates the correlation coefficient (R) between expression of the Wnt pathway gene and the indicated gene set. The upper panel shows the inverse association of MES (orange) and ADRN (blue) signatures to Wnt genes, while the lower panels show the correlations of the individual Wnt metagenes to the same pathway genes. Note the very similar pattern of correlation of WMG-1 (yellow) and MES signature to Wnt pathway genes and the inverse profile of WMG-2 (green) and WMG-3 (red) correlations. The bottom panel shows the correlation of the same genes with prognosis, with the y-axis representing Bonferroni corrected p-values of a Kaplan scan analysis using eventfree survival. Green bars indicate the correlation of high expression to good prognosis, while the red ones to poor prognosis. **(C)** WMG-1 expression in adrenergic, mesenchymal and neural crest cells (Versteeg/Etchevers Mixed NB 34). **(D)** Correlation between MES signature and WMG-1 in SEQC (GSE62564). Each point represents a tumour. **(E)** High expression of MES signature genes is correlated to good prognosis in the same data set.

In order to explore if Wnt may regulate NB lineage identity, we studied the relationship between individual Wnt pathway members and MES and ADRN lineage identity signatures in a primary NB dataset (SEQC498). MES and ADRN metagenes were constructed on the basis of the study of van Groningen et al (van Groningen et al. 2017). We then plotted the correlation of expression (correlation co-efficient, R) of each Wnt pathway member gene in relation to the expression of MES and ADRN signatures. In addition to the components of the Wnt pathway described in KEGG, we also included LGR receptors and their ligands, R-spondins. Wnt components showed a strong correlation to the expression of MES signature along a gradient, with R values ranging from 0.81 to -0.381 (Figure 3B, top panel). The association to the ADRN signature was also pronounced (R:-0.54 to 0.48), but interestingly displaying an inverse pattern to that of MES. The functional significance of this pattern could be underpinned, for example, by *FZD3*, which has been shown to be required for maintenance of dividing sympathetic neuroblasts and correlates with ADRN identity (Armstrong et al. 2011). In contrast, *FZD1* and *FZD6*, previously reported to be associated with chemoresistance and tumorigenic stem-cells respectively (Flahaut et al. 2009; Cantilena et al. 2011) correlate highly with MES identity (Figure 3B). Although the selectivity of Wnt-Frizzled interactions is poorly understood (Grainger and Willert 2018), our analysis suggests that Wnt responses and opposing functional outcomes in NC and NB may be regulated by Wnt receptor configurations.

We previously demonstrated that Wnt can drive the expression of diverse sets of genes in NB cells that display distinct co-expression patterns in primary tumours, which we represented with Wnt metagenes. We reasoned that differential regulation of Wnt metagenes in NB tumours could be due to the configuration of the Wnt pathway, i.e. the differential expression of genes encoding for the pathway components. We, therefore, assessed the correlation of expression between Wnt pathway members and our Wnt metagenes, WMG-1, 2, 3 and 4 (Figure 3B), which revealed a striking similarity of WMG-1 (yellow) with MES identity, and an anticorrelation with ADRN identity. This was also partially true for WMG-2 (green), but to a much lesser extent, whereas WMG-3 (red), which we had previously associated by high-risk and MYC signatures, was generally low in MES and high in ADRN. WMG-2 and 3 had inverse correlation profiles, consistent with their contrasting expression patterns in NB tumours. WMG-4 (grey) correlation patterns were not distinct.

Wnt pathway members also correlated with prognosis (Figure 3B bottom). High expression of genes with moderate correlation to MES or ADRN signature and high correlation to WMG-2 were the strongest indicators of good prognosis. Conversely, Wnt pathway genes correlating with ADRN signature were usually indicators of poor prognosis.

Focussing on WMG-1, we observed it was also significantly higher in NCC and MES cells lines compared to ADRN cells (Figure 3C), and in the SEQC dataset WMG-1 and the MES signature showed a remarkable correlation (R=0.927, p=9.0*10^-214^) (Figure 3D). This was also apparent in a second NB dataset containing 649 tumours (Supplementary Figure 1A). We had previously shown that WMG-1 expression in NB gene expression datasets correlated with intermediate risk NB, and in accordance with that, we found that the high expression of the MES signature metagene correlated with a better prognosis in the SEQC (Figure 3E) and also in the Kocak dataset of 649 tumours (Supplementary Figure 1B). As both our WMG-1 (Szemes et al. 2018) and MES (van Groningen et al. 2017) signatures are associated with Hippo-YAP/TAZ, it may be inferred that Hippo signalling, which has been implicated in NB tumorigenesis (Wang et al. 2015; Yang et al. 2017), may, at least at diagnosis, not be integral to high-risk neuroblastoma. Interestingly, the role(s) of Hippo signalling in NC development are not well established, but the observations made here and by others are consistent with a report showing the importance of Hippo-YAP/TAZ in neural crest migration (Hindley et al. 2016).

The transcription factor PRRX1 was pinpointed as being involved in MES/NCC-like identity by both recent landmark studies on NB heterogeneity (van Groningen et al. 2017; Boeva et al. 2017). Indeed the first of these directly demonstrated conversion of ADRN to MES lineages by *PRRX1* induction in SK-N-BE(2)-C cells. As we had reported Wnt3a/Rspo2 treatment leading to EMT (Szemes et al. 2018), we investigated the possible links between Wnt signalling and PRRX1. As shown in Figure 4A, *PRRX1* transcripts were significantly higher in NB cluster 2 (intermediate risk, see Figure 2). Analysing gene expression changes accompanying *PRRX1* induction in SK-N-BE(2)-C cells, we found that WMG-1 genes, high expression of which is characteristic of intermediate risk NB (Szemes et al. 2018), showed increased expression following 144 hrs of *PRRX1* induction, with the exceptions of *MSX2* and *BMP4*, which were increased at earlier timepoints (24-72 hrs). Induction of WMG-2 was also increased, although in a less uniform way, whereas WMG-3 and WMG-4 genes appeared to be partitioned in their response, with two subsets of genes regulated in an opposite fashion (Figure 4B). In sum, we found a remarkable overlap between our Wnt targets and the long-term transcriptional effects of *PRRX1* induction in SK-N-BE(2)-C. We conducted the same analysis on Wnt pathway genes (KEGG and other known members) and observed increased expression of Wnt pathway receptor and transcriptional regulator genes accompanying *PRRX1* induction (Supplementary Figure 2).

**Figure 4.**
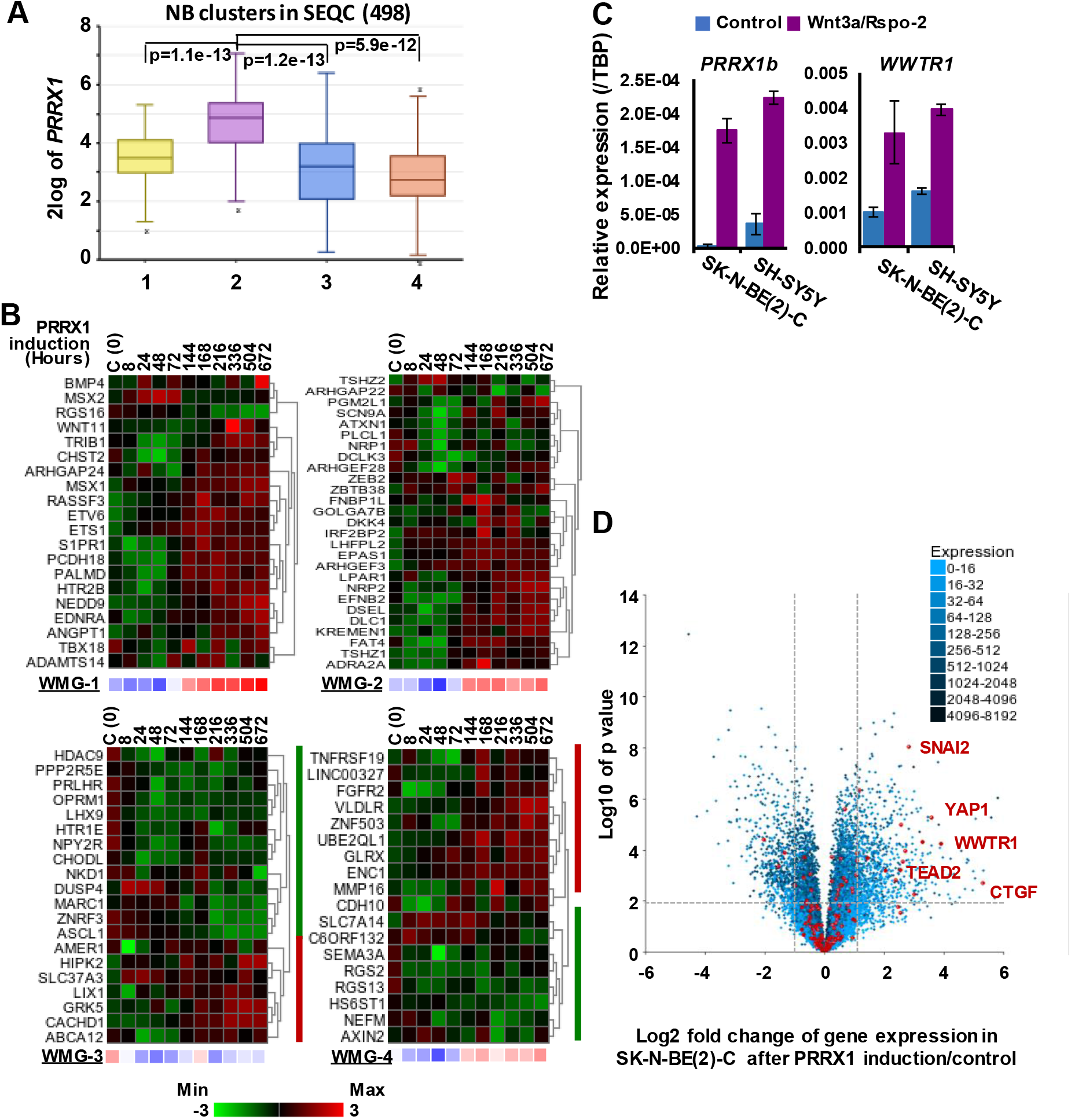
Associations between neuroblastoma Wnt target genes, *PRRX1*, and Hippo-YAP/TAZ signalling. **(A)** *PRRX1* expression is significantly higher in the intermediate risk NB Wnt cluster, which strongly associates with high WMG-1 expression (see Figure 1) (GSE62564). **(B)** Heatmaps showing the regulation of individual genes within the Wnt metagenes by PRRX1 (GSE908040, derived from inducible PRRX1 expression in SK-N-BE(2)-C cells). Note induction of WMGs 1, 2, and 4 at later timepoints, and downregulation of most WMG-3 genes. **(C)**. Wnt3a/Rspo-2 treatment of NB cell lines for 96 hours lead to the activation of MES-related transcription factors, *PRRX1*, and *WWTR1*. **(D)** Volcano plot showing gene expression changes in SK-N-BE(2)-C cells with induced PRRX1 expression. Red dots indicate members of the Hippo signalling pathway (KEGG). The highlighted Hippo pathway transcription factors, and the Hippo target gene *CTGF* were all strongly upregulated. Dashed lines separate genes regulated by minimum +/-2-fold with p<0.01 significance. Cell culturing and gene expression analysis were performed as previously (Szemes et al. 2018). PCR primers are available on request.

Although the kinetics of WMG responses argues against our Wnt genes being direct targets of PRRX1, this analysis links PRRX1, Wnt signalling and MES identity. We note that PRRX1 has been reported to be upstream of Wnt signalling and regulating EMT in gastric cancer (Guo et al. 2015). We also investigated whether Wnt signalling might form part of the regulatory circuit involving PRRX1, and the Hippo regulator TAZ (encoded by *WWTR1*), which were both implicated in MES/NCC-like identity. As these were not identified in our RNA sequencing timepoint (6 hrs), we assessed 96 hr treatments of SK-N-BE(2)-C and SH-SY5Y cell-lines by qPCR. Both *PRRX1* and *WWTR1* were upregulated by greater than 2-fold in both cell-lines (Figure 4C), suggesting indirect regulation by Wnt signalling. Finally, as we found strong interrelationships between WMG-1 with Hippo-YAP/TAZ, and PRRX1, we tested the possible link between PRRX1 and Hippo signalling. Evaluation of the *PRRX1* induction gene expression dataset revealed that numerous genes encoding for members of the Hippo signalling pathway were markedly upregulated, especially the Hippo effectors *YAP1*, *WWTR1*, their interacting transcription factors, *SNAI2* and *TEAD2*, and the archetypal YAP/TAZ target gene *CTGF* (Figure 4D). Taken together, this analysis supports Wnt signalling as a key regulator of the MES/NCC-like identity evident in NB.

Wnt signalling is known to crosstalk with Hippo-YAP/TAZ signalling, and an alternative Wnt pathway has been characterised showing that a non-canonical Wnt-YAP/TAZ signalling can be initiated by Wnt3a or Wnt5a/b, which can then suppress the canonical Wnt/β-catenin pathway (Park et al. 2015). Although Wnt3a can activate both canonical and non-canonical Wnt pathways (Samarzija et al. 2009), the majority of the genes we identified in SK-N-BE(2)-C cells had been shown to have β-catenin binding to their promoters (Szemes et al. 2018). Furthermore, we did not see early effects on YAP and TAZ proteins, or on *CTGF* expression (unpublished data) of Wnt3a/Rspo2 treatment in NB cells. Whilst this supports the early transcriptional response of SK-N-BE(2)-C cells to Wnt3a/Rspo2 to be canonical Wnt/β-catenin signalling, we also note that the non-canonical Wnt ligand genes, *WNT5B* and *WNT11*, were amongst genes upregulated by our treatments (Table 1). Thus, although it currently remains unclear what the relative contributions of canonical and non-canonical Wnt signalling are to NB cellular heterogeneity, it is evident that canonical Wnt signalling is interacting with other trans regulators to determine and maintain MES identity.

Our Wnt-driven NB phenotypes and transcriptome also indicated neural differentiation in SK-N-BE(2)-C and SH-SY5Y cell-lines, including induction of *NEFL* and *NEFM*. In terms of NB cellular identities, this suggests that Wnt may also influence ADRN (or noradrenergic) cell fates. As a first step to analysing this possibility, we determined the expression pattern of the ADRN signature genes in the four prognostic NB clusters defined by the expression of our NB Wnt target genes (Szemes et al. 2018) (Supplementary figure 3). ADRN genes had low expression in NB cluster 2, dominated by the overexpression of MES-related WMG-1. We also observed that the ADRN genes clustered into two subsets according to expression in the different prognostic Wnt groups. Therefore, we repeated the clustering in the SEQC dataset according to clinical risk categories and defined high-risk (ADRN-HR) and low-risk ADRN (ADRN-LR) signatures, based on high expression of the subsets in risk groups (Figure 5A). Contrasting expression of these two ADRN subsets was particularly striking in MNA tumours, which showed the highest expression of ADRN-HR and the lowest of ADRN-LR genes. Kaplan-Meier survival analyses showed that the ADRN-HR and ADRN-LR signatures were, respectively, strong negative and positive prognostic indicators (Figure 5B). We also found that ADRN-LR was positively correlated with WMG-2, which in turn negatively correlated with ADRN-HR. Evaluation of Hallmark genesets highlighted a striking positive correlation of ADRN-LR with Hedgehog signalling, as we had previously pointed out for WMG-2 (Szemes et al. 2018) (Figure 5C, top). The ADRN-HR signature negatively correlated with WMG-1, but positively with WMG-3, and accordingly, showed a very strong correlation with a MYC signature (Figure 5C, bottom). Gene Ontology analysis for ADRN-LR genes revealed enrichment for neuronal development and differentiation pathways, and in contrast, ADRN-HR genes were largely involved in cell-cycle processes, proliferation (Table 2). In keeping with the strong association between ADRN-HR and WMG-3 signatures, WMG-3 genes were expressed at significantly higher levels in ADRN cells compared to MES and NCC lines. WMG-4 displayed a similar expression pattern, albeit with lower significance. (Figure 5D). WMG-2 was not significantly different between MES and ADRN cell lines but was slightly lower in ADRN lines relative to NCC lines.

**Figure 5.**
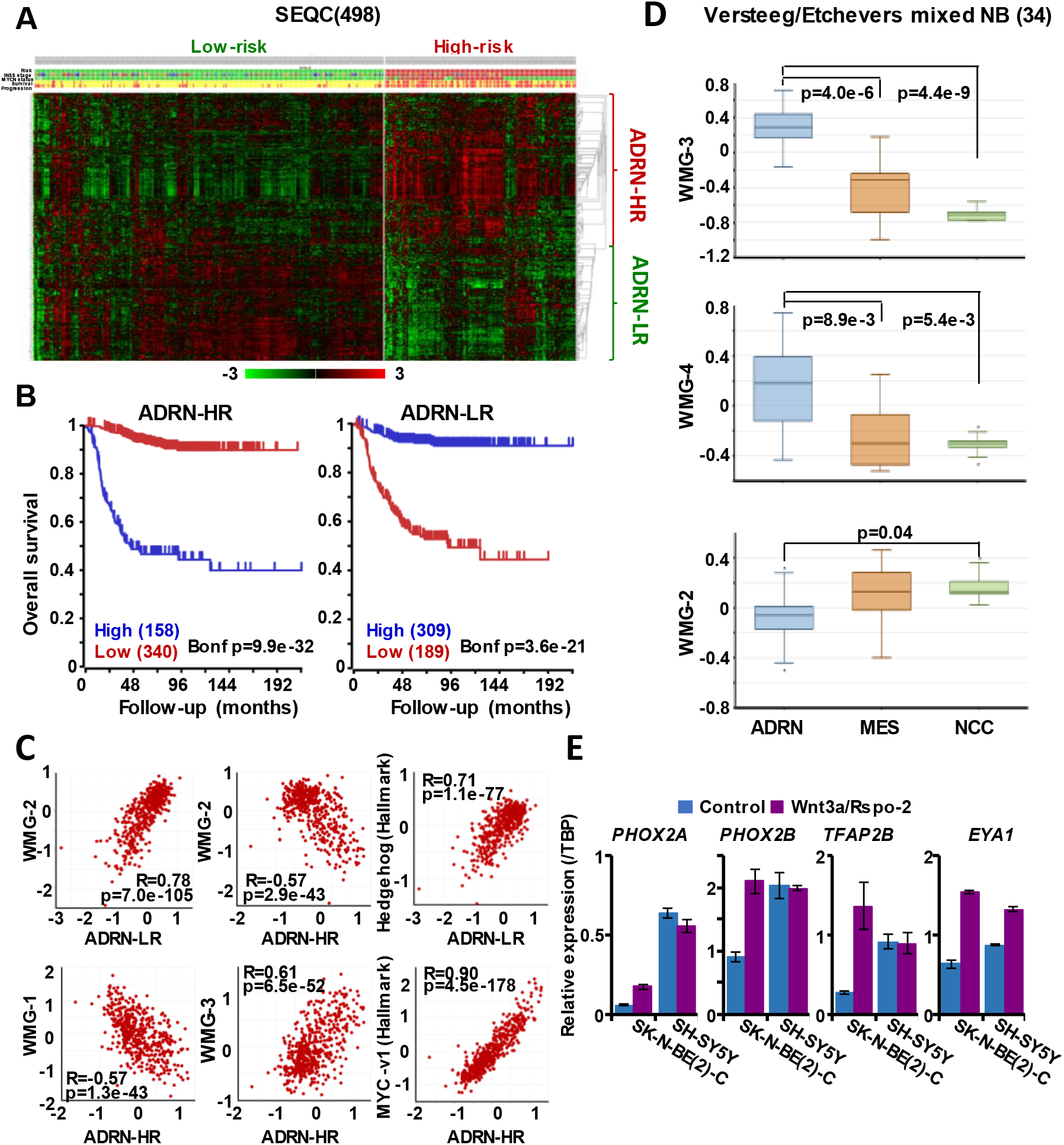
The ADRN signature can be partitioned and the subgroups correlate with prognosis and neuroblastoma Wnt-regulated metagenes. **(A)** Heatmap showing the expression of ADRN signature genes in risk groups of SEQC NB set (GSE62564) reveals two subgroups. **(B)** Kaplan-Meier analysis demonstrating that expression of the newly defined ADRN-LR and HR subgroups strongly and inversely correlate with prognosis. **(C)** WMG-2 shows a strong positive correlation with ADRN-LR, and negatively correlates with ADRN-HR. ADRN-LR has a positive association with Hedgehog signalling (Hallmark). ADRN-HR has a strong inverse correlation to WMG-1, and a positive correlation with WMG-3 and the expression of MYC targets-v1 (Hallmark). **(D)** WMG-3 and 4 are highly expressed in ADRN cells relative to MES and NC cells, and WMG-2 is significantly higher in NC cells than ADRN cells; WMG-2 is also higher in MES compared to ADRN, but the difference is not significant (Versteeg/Etchevers Mixed NB 34). **(E)** Long-term (96h) Wnt3a/Rspo-2 treatment of SK-N-BE(2)-C cell can activate ADRN lineage transcription factors in NB cell lines.

**Table 2.** Gene Ontology analysis of ADRN-LR and HR subgroups of genes.

| ADRN-LR |  |  |  |
| --- | --- | --- | --- |
| GO ID | Pathway description | N | FDR |
| GO.0007399 | nervous system development | 52 | 6.16E-12 |
| GO.0030182 | neuron differentiation | 37 | 6.16E-12 |
| GO.0048699 | generation of neurons | 43 | 6.16E-12 |
| GO.0022008 | neurogenesis | 42 | 7.11E-11 |
| GO.0048666 | neuron development | 32 | 8.62E-11 |
| GO.0031175 | neuron projection development | 29 | 1.45E-10 |
| GO.0048667 | cell morphogenesis involved in neuron differentiation | 24 | 1.30E-08 |
| GO.0048731 | system development | 64 | 1.30E-08 |
| GO.0048812 | neuron projection morphogenesis | 24 | 1.69E-08 |
| GO.0000904 | cell morphogenesis involved in differentiation | 26 | 2.12E-08 |
| ADRN-HR |  |  |  |
| GO ID | Pathway description | N | FDR |
| GO.0000278 | mitotic cell cycle | 28 | 2.41E-06 |
| GO.1903047 | mitotic cell cycle process | 26 | 2.72E-06 |
| GO.0022402 | cell cycle process | 30 | 6.23E-06 |
| GO.0007067 | mitotic nuclear division | 17 | 1.77E-05 |
| GO.0000280 | nuclear division | 18 | 5.86E-05 |
| GO.0007049 | cell cycle | 32 | 5.86E-05 |
| GO.0044772 | mitotic cell cycle phase transition | 14 | 0.00085 |
| GO.0006268 | DNA unwinding involved in DNA replication | 4 | 0.00161 |
| GO.0006271 | DNA strand elongation involved in DNA replication | 5 | 0.0104 |
| GO.0051301 | cell division | 15 | 0.012 |

Since we previously demonstrated that Wnt3a/Rspo2 treatment can prompt neural differentiation of NB cell-lines, we evaluated whether long-term (96 hr) treatments with Wnt ligands may influence transcription factors that contribute to the establishment of the ADRN, or noradrenergic, cell lineage. As shown in Figure 5E, *PHOX2A/2B, TFAP2B* and *EYA1* were all markedly upregulated in SK-N-BE(2)-C cells. In SH-SY5Y cells, these factors were already generally higher, but again induction of *EYA1* was apparent following Wnt3a/Rspo2 treatment.

Taken together these analyses demonstrate that the ADRN signature can be partitioned into two discrete subsets, which we have termed ADRN-LR and ADRN-HR, with contrasting expression in low-risk and high-risk NBs. The role of Wnt in differentiation of sympathoadrenergic neurons is further underlined by a report by Bodmer et al that Wnt5a promotes axonal growth and branching in response to NGF:TrkA signalling (Bodmer et al. 2009). Our data also suggest co-operativity of Wnt with Hedgehog signalling in establishing the ADRN-LR signature. The role of Hedgehog signalling in NB remains poorly understood and, as in the case of Wnt signalling, there is conflicting evidence; two studies have suggested its oncogenic potential (Mao et al. 2009; Xu et al. 2012), whereas others have reported an influence on differentiation of NB cells (Williams et al. 2000; Souzaki et al. 2010). A role for Hedgehog signalling in cephalic and trunk neural crest cell differentiation has been reported (Calloni et al. 2007), and in Xenopus, the Hedgehog transcription factor Gli2 was shown to be necessary for NCC specification and migration (Cerrizuela et al. 2018). In the context of NB, our previous studies suggested that WMG-2 genes were correlated with Hedgehog signalling and are suppressed by MYCN (Szemes et al. 2018). It therefore follows that ADRN-LR genes, strongly associated with neural development, may be Wnt/Hedgehog regulated genes repressed by MYCN. This informs and extends the suggestion that MYCN represses signalling pathways driving differentiation during NB tumorigenesis (Duffy et al. 2015).

### Summary & Perspectives

The understanding of gene expression programmes, contingent on transcription factor hierarchies and chromatin level plasticity, in particular epigenetic changes at super-enhancer elements, continues to facilitate our understanding of cellular identity. Notable advances have been made in ascertaining regulatory circuits in neural crest cells as well as in neuroblastoma recently, yet much remains to be determined about the contributions of autocrine and paracrine signalling pathways that, together with transcription factors and epigenetic states, forms a regulatory triumvirate.

Our analysis of Wnt signalling strongly suggests that this pathway is a major determinant of transcription factor circuitries, and thereby lineage identity, in neuroblastoma. This is further supported by comparisons with NC GRNs, with Wnt signalling regulating early NC specifiers such as *MSX1* and *DLX5/6*, as well as migratory factors such as *SNAI2* and *TWIST1/2*. Further, we highlight the possible interplay between canonical Wnt signalling with Hippo-YAP/TAZ and Hedgehog signalling, emphasizing the need for integrative analysis of signalling inputs for both NC and NB biology. Our transcriptomic signatures facilitate this, but also underline the need for parallel studies using diverse cell types and signalling effectors.

With regard to NB, our study begins to reconcile some of the apparent contradictions about the influence of Wnt signalling. The determination of MES/NCC-like identity by Wnt is consistent with reports that resistance to chemotherapy and stemness may be linked to Wnt (Flahaut et al. 2009; Cantilena et al. 2011) (Vangipuram, Buck, and Lyman 2012). However, as we have shown, there are also Wnt-driven gene modules linked to neuronal differentiation. We propose that establishing a “Signalling Regulatory Network” will be necessary to formulate novel and efficacious treatment combinations for resistant NB and differentiation therapies.

## Acknowledgements

We wish to thank Children with Cancer UK, the Childrens Cancer and Leukaemia Group (CCLG), Neuroblastoma UK and Smile with Siddy, the Biotechnology and Biological Sciences Research Council (BB/P008232/1) and Cancer Research UK (A12743/A21046) (to K.M.) for funding this study.

## Abbreviations

ADRN: adrenergic
EMT: epithelial-mesenchymal transition
ERK: extracellular signal-regulated kinase
GRN: gene regulatory network
KEGG: Kyoto encyclopedia of genes and genomes
LEF: lymphoid enhancer binding factor
MEK: MAPK/ERK kinase
MES: mesenchymal
MNA: *MYCN*-amplified
NB: neuroblastoma
NC: neural crest
NCC: neural crest cell
Rspo: R-spondin
SA: sympathoadrenal
TCF: T-cell-factor

## Notes

Declarations of interest: none

