## Supplementary Figures 1-3 for "Wnt signalling is a major determinant of neuroblastoma cell lineages"

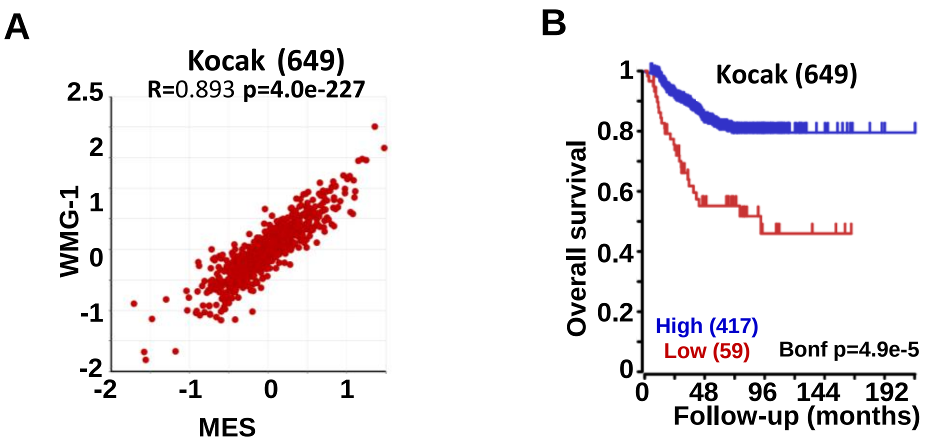


**Supplementary figure 1.**

**(A)** Correlation between MES signature and WMG-1 and **(B)** Kaplan-Meier survival analysis of MES genes in Kocak(649) data set (GSE45547).

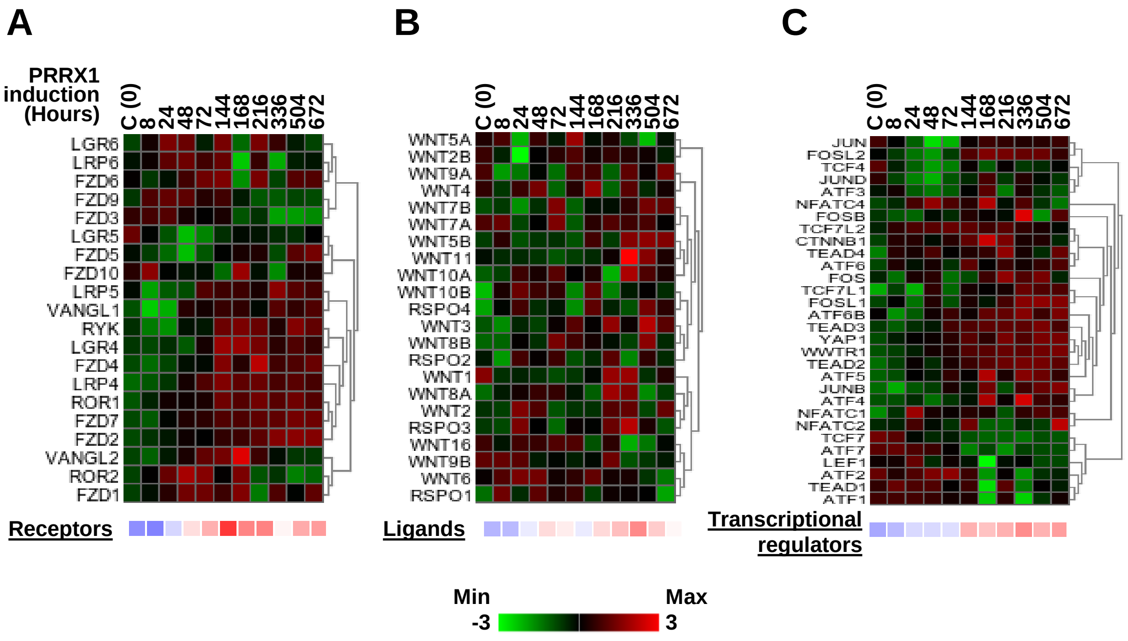


**Supplementary figure 2.**

Heatmaps showing the regulation of the Wnt receptors (A), ligands (B) and transcriptional regulators (C) in PRRX1-induced SK-N-BE(2)-C cells (GSE90804). In addition to KEGG-derived Wnt pathway members, we included other known regulators as well. *LGR4-6* were added to the receptor group and *Rspo-1-4* to the ligands **[14]**. Wnt-associated transcriptional regulators also included *YAP1*, *WWTR1*, *TEAD1-4*, *ATF1-7*, *FOS*, *FOSB*, *FOSL1/2*, *JUNB*, *JUND* and *TCF4* **[9, 11]**.

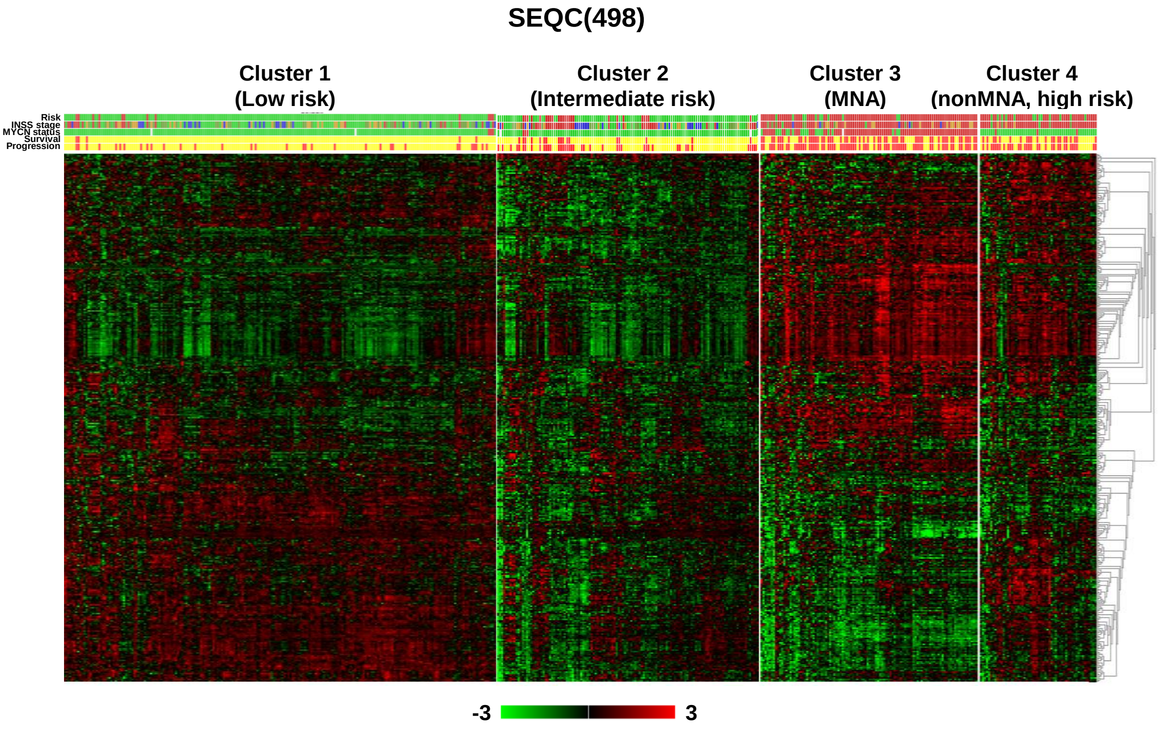


**Supplementary figure 3.**

Expression of ADRN signature genes in prognostic, Wnt-related NB clusters in SEQC data set (GSE62564) revealed two subgroups that correlated with risk status.
